# Frequency-dependent competition between strains imparts resilience to perturbations in a model of *Plasmodium falciparum* malaria transmission

**DOI:** 10.1101/2020.12.22.423749

**Authors:** Qixin He, Shai Pilosof, Kathryn E. Tiedje, Karen P. Day, Mercedes Pascual

## Abstract

In high-transmission endemic regions, local populations of *Plasmodium falciparum* exhibit vast diversity of the *var* genes encoding its major surface antigen, with each parasite comprising multiple copies from this diverse gene pool. This strategy to evade the immune system through large combinatorial antigenic diversity is common to other hyperdiverse pathogens. It underlies a series of fundamental epidemiological characteristics, including large reservoirs of transmission from high prevalence of asymptomatics and long-lasting infections. Previous theory has shown that negative frequency-dependent selection (NFDS) mediated by the acquisition of specific immunity by hosts structures the diversity of *var* gene repertoires (strains), in a pattern of limiting similarity that is both non-random and non-neutral. A combination of stochastic agent-based models and network analyses has enabled the development and testing of theory in these complex adaptive systems, where assembly of local parasite diversity occurs under frequency-dependent selection and large pools of variation. We show here the application of these approaches to theory comparing the resilience of the malaria transmission system to intervention when strain diversity is assembled under (competition-based) selection vs. a form of neutrality, where immunity depends only on the number but not the genetic identity of previous infections. The transmission system is considerably more resilient under NFDS, exhibiting a lower extinction probability despite comparable prevalence during intervention. We explain this pattern on the basis of the structure of strain diversity, in particular the more pronounced fraction of highly dissimilar parasites. For simulations that survive intervention, prevalence under specific immunity is lower than under neutrality, because the recovery of diversity is considerably slower than that of prevalence and decreased *var* gene diversity reduces parasite transmission. A Principal Component Analysis of network features describing parasite similarity reveals that despite lower overall diversity, NFDS is quickly restored after intervention constraining strain structure and maintaining patterns of limiting similarity important to parasite persistence. Given the resulting resilience to perturbations, intervention efforts will likely require longer times than the usual practice to eliminate *P. falciparum* populations. We discuss implications of our findings and potential analogies for ecological communities with non-neutral assembly processes involving frequency-dependence.

## 1 Introduction

One major goal of community ecology has been to demonstrate that system-level patterns are influenced by specific interactions and are not the mere consequence of stochastic birth-death processes, such as immigration, growth, and extinction (Hubbell, 2001). This problem lies at the heart of rejecting the neutral theory of community ecology. The presence of a particular ecological interaction at the ‘microscopic’ level of individuals is not sufficient and does not guarantee its importance at the ‘macroscopic’ level of populations or communities (Azaele *et al*., 2016). Less appreciated is the analogy between these lines of research and those in infectious disease ecology concerning pathogen antigenic diversity (Pascual, 2020). In community ecology, competition for resources and selection for traits that give an edge in resource acquisition shape the biodiversity of plants and animals and the structure of species interactions. In infectious diseases, competition for hosts and selection for traits that allow pathogens to evade human immune systems shape the antigenic diversity and underlying genetic population structure of pathogens (Gupta & Day, 1994; Grenfell *et al*., 2004; Koelle *et al*., 2006; Volz *et al*., 2013). Theory on the interface of such dynamics is found in the fields of phylodynamics and strain theory (Gupta & Day, 1994; Grenfell *et al*., 2004).

One system in which these analogies have been investigated is that of the malaria parasite *Plasmodium falciparum*. Strain structure and coexistence in local populations of the malaria parasite result from an on-going assembly process involving the dynamic interplay between ecology (population dynamics of strains, competition for hosts) and evolution (genetic changes via mutation and recombination) (He *et al*., 2018). The trait of interest concerns the variation in the major antigen of the blood stage of infection PfEMP1. This protein is exported by the parasite to the surface of the infected blood cells where it becomes the target of the adaptive immune system (Bull *et al*., 1998). Variations of the protein, or variable surface antigens (VSAs), are encoded by a multigene family known as *var* with about fifty to sixty gene copies per parasite, whose expression is sequential and influences the duration of infection. From this perspective, an individual parasite corresponds to a unique combination of *var* genes — a ‘repertoire’ — encoding for a particular set of VSAs. In high-transmission endemic regions such as those of sub-Saharan Africa, local diversity of *var* gene types is in the order of tens of thousands. This extremely large genetic pool underlies the vast combinatorial diversity of the repertoires themselves. Both these high levels of diversity result from high recombination rates, which act at the two different levels of organization, mixing repertoires and generating new *var* genes respectively (Zilversmit *et al*., 2013; Larremore *et al*., 2013).

Previous theory and data have shown that frequency-dependent competition for hosts is at play in determining the coexistence of a large number of repertoires, whose population structure is both non-random and non-neutral despite the vast gene pool (Barry *et al*., 2011; Day *et al*., 2017; He *et al*., 2018; Pilosof *et al*., 2019). Hosts can be viewed as resource patches whose availability depends on their individual history of ‘consumption’ from previous exposure. This is because exposure to a specific VSA leads to the acquisition of specific immunity by individual hosts, which then shortens infection reducing the fitness of the parasites that carry the corresponding gene. Adaptive immunity therefore provides an advantage to strains carrying rare *var* genes, and a disadvantage of those composed of common ones. In the language of community ecology, the system experiences stabilizing competition (sensu Chesson, 2000) from trait variation that underlies niche differences. In that of population genetics, such competition is a mechanism of negative frequency-dependent selection (NFDS) which acts as a form of balancing selection and promotes strain (i.e., *var* repertoire) coexistence (He *et al*., 2018; Pilosof *et al*., 2019).

We have previously used the malaria system to address the challenge of discerning rules of assembly from complex population patterns, focusing on the large strain variation of *P. falciparum* populations in regions of high transmission. We applied for this purpose a combination of stochastic agent-based models and network analyses of the genetic similarity between repertoires they generate (Artzy-Randrup *et al*., 2012; He *et al*., 2018; Pilosof *et al*., 2019). These studies have shown that networks assembled by NFDS have different topologies than those assembled in the absence of selection (i.e., under neutral processes). Hence, the malaria system teaches us that the importance of individual and specific interactions can be detected in features of the macroscopic similarity structure of (strain) diversity. We note that networks rather than phylogenetic trees were applied because of the predominant role of recombination in the evolutionary change of the system. Trees have been employed instead in pathogen phylodynamics when considering evolution via mutation (Grenfell *et al*., 2004; Koelle *et al*., 2006; Volz *et al*., 2013; Zinder *et al*., 2013). Regardless of the particular graphs utilized, their combination with stochastic ABMs proved fruitful and may find more general application to describe patterns of limiting similarity, identify the importance of specific ecological processes, and test them against appropriate neutral counterparts.

The structure of diversity has been a major driver of theory development in community ecology, because of its hypothesized influence on system resilience to perturbations and therefore species’ extinction (Hutchinson, 1959; Allesina & Tang, 2015). In infectious disease dynamics, a parallel yet little studied line of research regards the resilience of structured pathogen populations to interventions. Here, we illustrate the combination of stochastic ABMs and network analysis for addressing the resilience of *P. falciparum* populations to interventions that decrease transmission, depending on neutral vs. non-neutral assembly. We ask whether the malaria system assembled under NFDS is more or less resilient to perturbations than a neutral counterpart. We apply press perturbations (Ives & Carpenter, 2007) in a stochastic computational model to represent interventions that target the mosquito vector with indoor residual spraying (IRS), which involves the application of an insecticide to internal walls and ceilings of homes (WHO, 2015). IRS effectively reduces transmission rate and therefore, the growth rate of the parasite population as a whole. We consider resilience in terms of the persistence of parasite populations across intervention, as well as the recovery of prevalence (abundance) and genetic diversity post-intervention. We then consider the response of repertoire structure as a manifestation of whether the influence of NFDS recovers quickly after the intervention. We end by drawing plausible analogies and implications for other ecological systems (Forrister *et al*., 2019a; Azarian *et al*., 2019), in particular for communities of species under stabilizing competition, Janzen-Connell mechanisms of coexistence and other forms of balancing selection.

## 2 Materials and Methods

### 2.1 Agent-based model (ABM) and model setup

Malaria transmission and intervention are modeled using an agent-based, discrete-event, continuous-time stochastic system described in detail in He *et al*. (2018) and Pilosof *et al*. (2019). Here, we briefly describe the ABM, with an emphasis on the specific implementation of the regional intervention scheme that constitutes the press perturbation.

We model a local parasite population, *N*, as well as a global parasite pool, *G*_*p*_ that acts as a proxy for regional parasite diversity (Table **??**). Parasite genomes can migrate from the regional pool to the local population. Each simulation starts with 20 migrant infections to initiate local transmission (Fig. 1A). Because each parasite genome is a repertoire of 60 *var* genes, migrant genomes are assembled from random sampling of 60 *var* genes from the global pool. Each *var* gene is itself a linear combination of two epitopes – the part of a molecule that acts as an antigen and is recognized by the immune system (Rask *et al*., 2010; He *et al*., 2018) (Table **??**).

**Figure 1:**
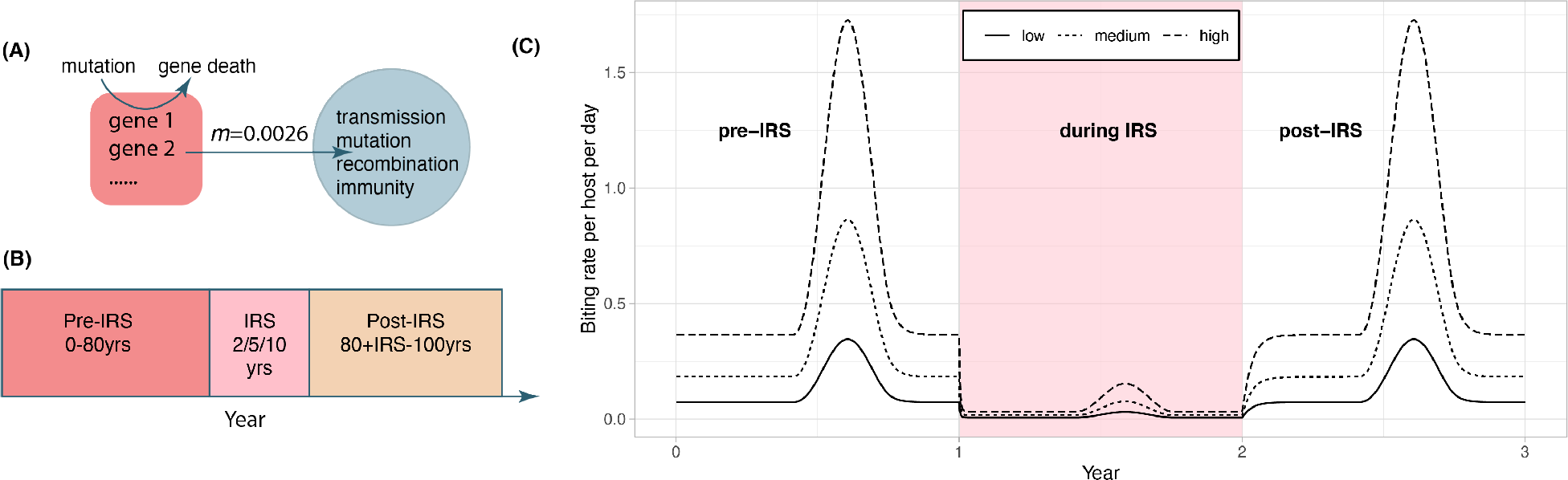
Schematic of the IRS experiment design. (**A**) Initial transmission events and migrant genomes in a local population (blue circle) are sourced from a global pool of genes (red square). Mutation of new genes and the death of existing genes in the global pool occur at the same rate as in the local population (see Methods). The value of the migration rate is inferred from an empirical dataset for a high transmission region in Ghana (see Methods). Individual infections are tracked locally, as well as events for transmission, mutation of genes, recombinatios within and between genomes, and acquisition and loss of specific immune memory in hosts. (**B**) Each simulation follows three stages (after a burn-in period): a pre-IRS period during which the transmission in the local population reaches a stable state; an IRS period of 2, 5, or 10 years reducing transmission, and a post-IRS period when transmission rates go back to pre-IRS levels. (**C**) Three levels of transmission intensity (biting rates) are explored in the experiments (pre-/post-IRS, low: 44 bites per host per year; medium: 110 bites/h/y; high: 221 bites/h/y).

We consider seasonal endemic transmission dynamics in which mosquitoes are not represented explicitly (i.e., as agents in the model), but via biting events. Local transmission events are sampled at the total rate of host population size *N*_*host*_ times the biting rate *b*, in which a donor and a recipient host are selected randomly. We implemented seasonality as fluctuation of mosquito bites via density dependence at the egg and larva stages as a function of rainfall typical of Northern Ghana (availability of breeding sites White *et al*., 2011). The specific algorithm to obtain the seasonality vector was described in detail in Pilosof *et al*. (2019).

The main modification to the current model is how the global pool interacts with local transmission. First, instead of remaining static, the global gene pool in this implementation updates its gene composition at the same mutation rate as that of the local population. Specifically, new genes are generated at the rate of the product of local parasite population size and the per-allele mutation rate. Once a new gene is generated, the old gene that it mutates from is removed from the global gene pool. Genomes migrate from the global pool to the local population (Fig. 1A). The number of migrant genomes increases in wet seasons and decreases in dry seasons (see details in section Estimation of migration rate). Interventions are assumed to be applied at the regional level so that prevalence of the disease is the same across the region, including the local population. Therefore the proportion of infective bites from migration is kept the same as that of local transmission.

### 2.2 Course of a simulation and Indoor Residual Spraying (IRS) Intervention

Each simulation follows three stages (Fig. 1B): (i) a pre-IRS period (0-80 yr) during which the local parasite community is assembled and the transmission system reaches a stationary state before the perturbation; (ii) an IRS period of 2, 5, or 10 years during which transmission is decreased; and (iii) a post-IRS period when transmission rates return to pre-IRS levels and the system is allowed to recover (Fig. 1C). During the IRS interventions, the effectiveness of insecticides in killing adult female mosquitoes is set to be 100% (Wanjala *et al*., 2015) and the percentage of sprayed household is set to be 90% (WHO, 2015; Coleman *et al*., 2017; West *et al*., 2014; Kigozi *et al*., 2012). Details on the model can be found in White *et al*. (2011), and our implementation (in Mathematica) at the GitHub repository https://github.com/pascualgroup/Pf_temporal_networks.

### 2.3 Within-host dynamics

The infection and immune history of each host are tracked individually. Upon each biting event, if the donor harbors parasites, then the strain has a given probability, *c*, to be transmitted to the mosquito. If the host harbors *m* infections concurrently, each parasite strain experiences a reduced transmissibility of 0.5*/m*. Var repertoires picked up by the mosquito at a bite event can recombine with another genome to produce sporozoites. Specifically, if there are *n* parasite genomes, each genome has a probability 1*/n* of recombining with itself, producing the same offspring genome, and a probability 1 *−* 1*/n* of recombining with a different genome, producing recombinants. This sexual recombination process creates variation at the genome level. The total number of *var* repertoires passed onto the receiving host is kept the same as that obtained from biting the infectious donor host.

Once in the blood, each infection progresses with the sequential expression of the *var* genes in random order. During an infection, novelty at the gene level can be generated via mutation or recombination between *var* genes within the same genome. Part of the life cycle of *Plasmodium falciparum* occurs in the liver; we therefore implement a 14 day delay to mimic the transition from the liver stage to the blood stage, after which the *var* genes are expressed and immunity towards them is gained.

Hosts gain and lose immunity towards specific epitopes, which we implement with a counter. Upon transition to expression of a different *var* gene or clearance of a *var* gene by the immune system, the host gains specific immunity towards the epitopes in the gene that was previously expressed. For example, if the host was infected by a parasite that contained two *var* genes, each of which with epitope *x*, the counter for epitope *x* will increase by two. The counter reduces by one at the immune loss rate of 1/1000 per day (Collins *et al*., 1968). When the number becomes 0, the host loses protection against the epitope.

### 2.4 Estimation of migration rate

Diversity patterns of microsatellite markers often indicate adjacent populations to be panmictic (Rorick *et al*., 2018). Here, we use empirical data of *var* genes, hyper-diverse markers that provide a higher resolution for recent migration events, to infer the rate of gene exchange between populations and to obtain a reasonable estimate for migration rates that allows us to implement an open transmission system. We used Jost’s *D* measure of population divergence (Eq. 12 in Jost (2008)) to consider highly diverse genetic markers, and to compare *var* gene composition within and between two field sites for which molecular sequences were previously obtained, Soe and Vea/Gowrie in the Bongo District of Ghana at the end of the wet or high transmission season (He *et al*., 2018; Tiedje *et al*., 2017). Using Eq. 22 in Jost (2008), we estimated the migration rate by dividing *D* by the mutation/recombination rates of the *var* genes per generation (5.3e-6, Claessens *et al*. (2014)).

### 2.5 Event scheduling in the stochastic model

The simulation is an agent-based model implemented with a Gillespie algorithm, which is a computationally-efficient way to model stochastic dynamics. The algorithm takes a set of events (e.g., infection) and randomly chooses the next event and the time in which it will occur. The probability of choosing the next event is proportional to the abundance of entities that compose it. For example, the probability of infection with a given repertoire depends on its abundance. The time interval to the next event is randomly chosen from an exponential distribution with a mean of 1 over the sum of rates. In our simulation, global events include local transmission from biting events, new transmission from migrant *var* repertoires, and birth and death of hosts. In the numerical implementation of the simulation, all possible future events are stored in a single event queue along with their putative times. When an event occurs, it may trigger the addition or removal of future events on the queue, or changes of their rates, leading to a recalculation of their putative time. This implementation is adapted from the next-reaction method following Gibson & Bruck (2000), which optimizes the Gillespie first-reaction method (Gillespie, 1976) and allows for faster simulation times as it targets changes of rate events to a given subset (specified by the structure of the queue).

### 2.6 Experimental design and selection regimes

We explored how initial gene pool size, transmission rate and the duration of an IRS intervention influence the system’s resilience to intervention. Specifically, we set: (i) initial gene pool sizes of 1200, 1800, 2400, 3600, 4800, 7200, 9600, 14400, and 19200; (ii) three levels of transmission intensity (biting rates): 44, 100, and 221 bites per host per year, corresponding to ‘low’, ‘medium’ and ‘high’ transmission (Fig. 1C); and (iii) IRS lasting for 2, 5, or 10 years. This design has 81 sets of parameter combinations and we ran 50 replications for each combination. We calculated the probability of extinction as the proportion of the 50 replications in which the parasite populations crashed before lifting the IRS.

Our main objective was to compare responses to perturbation in the presence and absence of frequency-dependent competition. Specifically, we aimed to compare an ‘immune selection scenario’, in which immune memory to particular epitopes elicits competition between parasite genomes for hosts, to a ‘neutral scenario’ of “generalized” immunity. In both these scenarios, duration of infection is the relevant trait. Parasite fitness is affected by immunity in our model through the duration of infection, since shorter duration reduces the rate of transmission. The duration of infection in a host depends on the host immune history of given *var* epitopes, as described above. This is the process that implements an advantage to rare genes and the parasites that carry them, and a disadvantage to common ones – that is, the frequency-dependent selection. The model of generalized immunity retains protection conferred by previous infection but does not consider specific memory towards *var* genes. This is a common implicit assumption of most malaria transmission models, including other ABMs and extensions of the well-known compartmental population models of the SIR type (for Susceptible-Infected-Recovered classes), where infection ‘comsumes’ susceptibles but as a general and shared resource. In our implementation of generalized immunity, the duration of infection depends on the number of previous infections but not on their specific genetic composition. For a meaningful comparison to the immune selection scenario, we parameterized the function of infection duration with number of previous infections, to match the curve generated by the corresponding immune selection scenario (see He *et al*., 2018). We refer hereafter to the scenario under immune selection or frequency-dependent selection as “S” and to that under generalized immunity as “G”. Comparisons are always made for the same parameter combinations.

### 2.7 Sampling and Gene Similarity Networks

During each simulation, summary statistics including prevalence, multiplicity of infection (MOI; number of genomes within a host), genetic and allelic diversity, were calculated every 30 days. In addition, 100 infected hosts were sampled to analyze parasite diversity patterns. To evaluate similarity of parasites in the population, pairwise type sharing (PTS) was calculated between all repertoire pairs (regardless of the host in which they are encountered) as *PTS*_*ij*_ = 2*n*_*ij*_*/*(*n*_*i*_ + *n*_*j*_), where *n*_*i*_ and *n*_*j*_ are the number of unique alleles (corresponding to epitopes) within each repertoire *i* and *j* and *n*_*ij*_ is the total number of alleles shared between repertoires *i* and *j* (Barry *et al*., 2007). In addition, similarity networks based on the *var* composition were built to investigate changes of parasite genetic population structure through interventions. We calculated genetic similarity of repertoire *i* to repertoire *j* as *S*_*ij*_ = (*N*_*i*_ *∩ N*_*j*_)*/N*_*i*_, where *N*_*i*_ and *N*_*j*_ are the number of alleles unique for repertoires *i* and *j*, respectively (the genetic similarity of repertoire *j* to repertoire *i* was calculated as *S*_*ji*_ = (*N*_*i*_ *∩ N*_*j*_)*/N*_*j*_). We used a directional index because of the asymmetric competition resulting from different numbers of unique alleles in a repertoire (He *et al*., 2018).

### 2.8 Calculation of Network properties and Principal Component Analysis

We calculated 36 network properties to compare the changes in network structure between the immune selection and neutral scenarios. These properties include metrics of transitivity, degree distributions, component sizes, diameters, reciprocity, and proportion of 3-node graph motifs (see Table S1 in (He *et al*., 2018) for a complete list of properties and definitions). In the network analyses we retained only edges with values within the 95% percentile to focus on the strongest interactions of current competition between strains (He *et al*., 2018; Pilosof *et al*., 2019). To minimize the influence of sample size differences across time due to changes in mean MOI, network properties were calculated by resampling 100 repertoires randomly from the original network. Principal component analysis (PCA) was performed on normalized and centered network properties across time and selection regimes per parameter combination, to inspect the overall trend of diversity structure.

## 3 Results

### 3.1 Populations under immune selection are more resilient

The interventions implemented in the model reduce the transmission rates by about 90% during IRS. Not surprisingly and especially for the long-lasting interventions, the stochastic simulations are prone to extinction. We computed the extinction probability from replication of the dynamics for a given parameter combination described in the Methods and found that a lower probability under specific immunity (S) than under generalized immunity (G) for the same parameters (Fig. 2A, Fig. S1A). As expected, longer interventions lead to higher extinction rates, especially under low transmission rates. A higher initial pool size of genetic diversity ensures a lower extinction probability. Pool sizes larger than 10,000 *var* genes, which represent high transmission endemic regimes in sub-Saharan Africa, experience significantly less extinction, even after 10 years of sustained intervention. Interestingly, this difference does not reflect a trivial effect of overall population size of the parasite: the mean prevalence is comparable during IRS for S and G, only slightly higher for the former, indicating that other factors are responsible for the higher resilience under NFDS (Fig. 2B, Fig. S1B).

**Figure 2:**
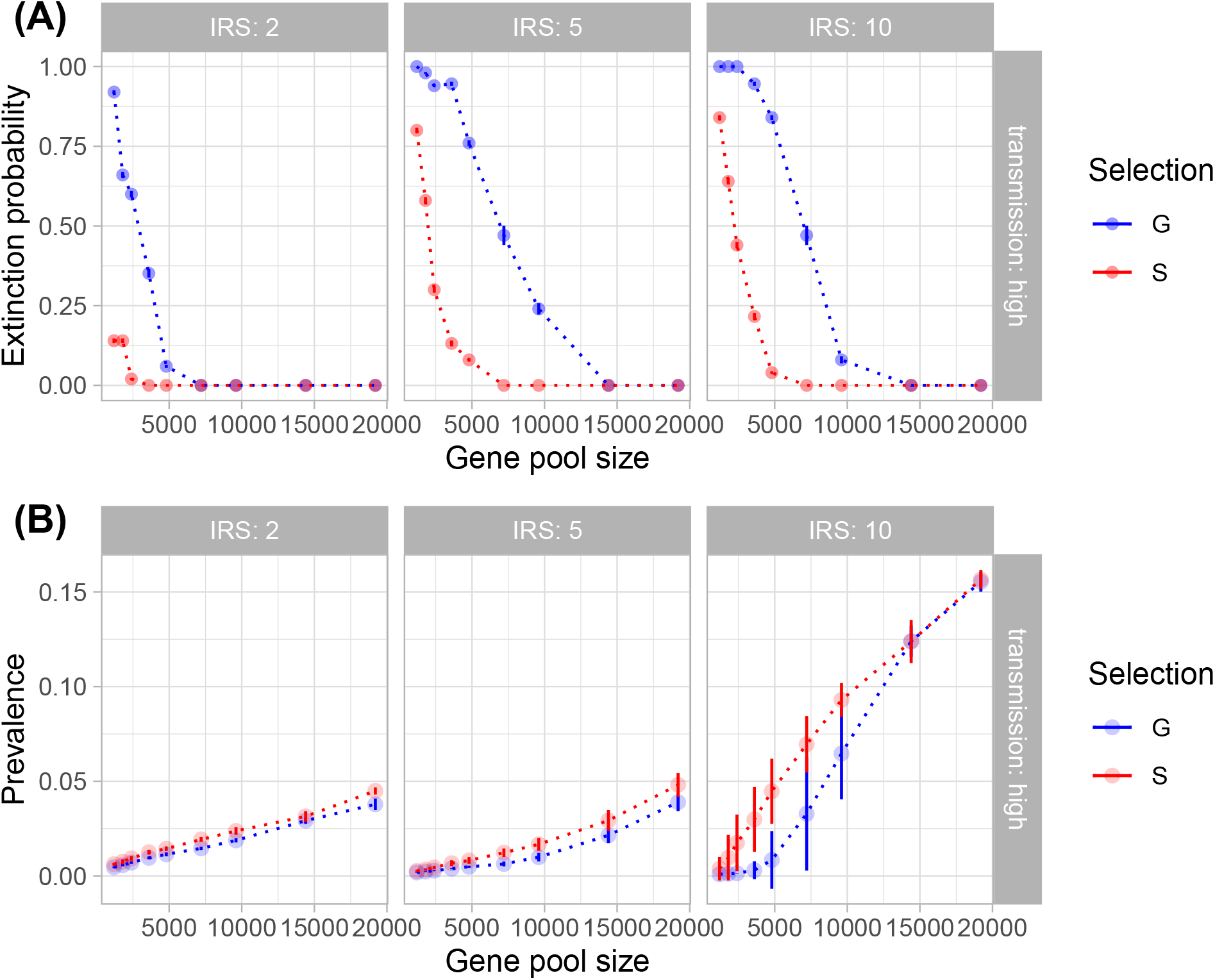
Extinction probability (A) and mean prevalence during IRS (B) as a function of initial gene pool size and lengths of IRS (columns). Shown here are the results from high transmission intensities. See Fig. S1 for results from low and medium transmission intensities. In (A), each point is the proportion of simulation runs for the given parameter combination (out of 50 runs) that crashed before the IRS was lifted. In (B), each point shows the mean value of 50 simulation runs for the given parameter combination, with bars representing the standard deviation. Blue color depicts simulations under generalized immunity (G, the neutral model), while red color corresponds to specific immunity (S, therefore NFDS). It is clear that despite comparable prevalence, the parasite population is less prone to extinction in the immune selection scenario.

### 3.2 Resilience is enhanced by persistent repertoire dissimilarity under NFDS

To investigate the origin of the difference in resilience as measured by the probability of extinction, we consider the epidemiological parameter that confers the fitness difference to the parasite, namely the duration of infection in the human host. As is typical in epidemiology, the reproductive number of the parasite is the product of the transmission rate and the duration of infection. In our model, immunity modifies only this duration as genes encoding for antigenic variants which have been seen before are not expressed. We note that we set the distribution of duration of infection with the number of previous exposures under G to exactly match that under S before IRS for a specific parameter combination. Therefore, the mean values of duration of infection under G and S are similar before IRS by construction, with the values increasing significantly during IRS (Fig. S2). When the longest infections are considered by comparing the top 5% of the distribution in infection duration, we observe higher values under S during IRS in less than a year (Fig. 3A, Fig. S3). Longer lifetimes of infection at the tail of the distribution provide a buffer against extinction by conferring a higher probability of surviving the low transmission period imposed by the intervention.

**Figure 3:**
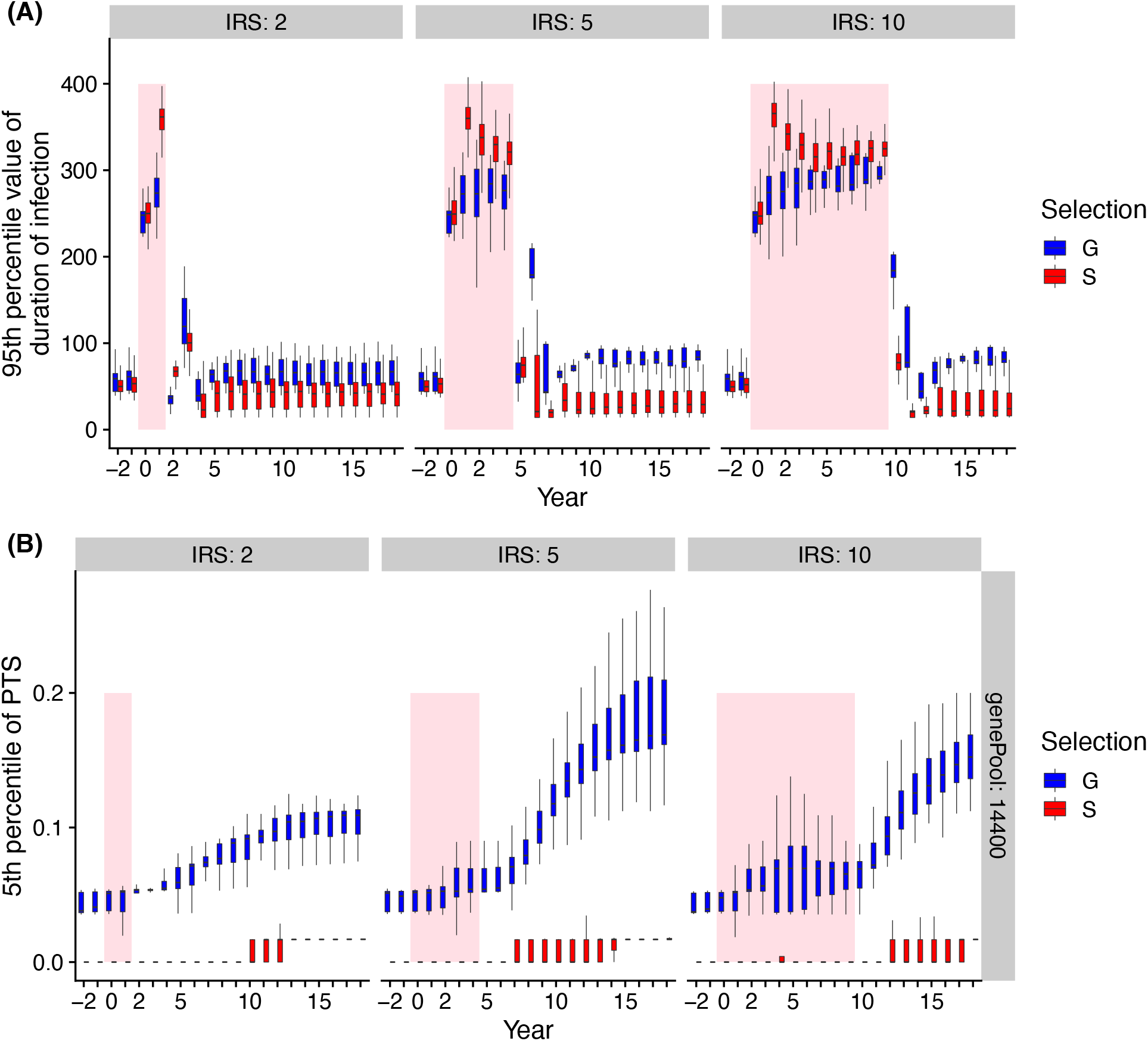
Comparisons of duration of infection and mean PTS over time under high transmission between different selection regimes. Boxplot summarizes the distribution of duration of infection in its 95th percentile (**A**) and 5th percentile of PTS between *var* repertoires (**B**) among replication runs. Shaded area indicates IRS period. Negative years correspond to years prior to IRS.

To further understand the extinction patterns, we therefore need to explain the longer duration of infection under NFDS. We turn to the structure of similarity between parasites as reflected by the distribution of pairwise type sharing or PTS between *var* repertoires (Fig 3B, Fig. 4, Fig. S4). Figure 4 shows the PTS distributions before, during and after the IRS intervention for the two regimes. Their shapes differ substantially at low PTS reflecting clear patterns of high dissimilarity. In particular, NFDS generates a monotonically decreasing distribution with the highest frequencies found at the lowest overlap. In contrast, under G the distribution exhibits a mode, with the highest frequency for some intermediate overlap. Due to decreased diversity during IRS, the proportion of repertoire pairs with high PTS increases for both scenarios, and the whole distribution moves towards higher overlap (Fig. 4). However, the change is less pronounced under S, especially for the maintenance of dissimilar repertoires (Fig. 3B). When the intervention is lifted, the distribution of G shifts significantly towards increased similarity, while the bimodal shape of PTS distribution in S is maintained, including the considerable fraction of highly dissmilar strains. Since diversity under G does not influence transmission by construction, change in PTS reflects solely parasite population fluctuations and bottleneck effects during and after IRS. The maintenance of low PTS values to pre-IRS levels under S is indicative of variant-specific immune selection at work: even under reduced diversity, repertoires are maintained as different from each other as possible, resulting in longer duration of infection in certain hosts than for infections with randomly assigned gene composition. A larger fraction of the parasite population with highly dissimilar repertoires generates higher heterogeneity in duration of infection, including longer duration, and strengthens resilience towards interventions under S.

**Figure 4:**
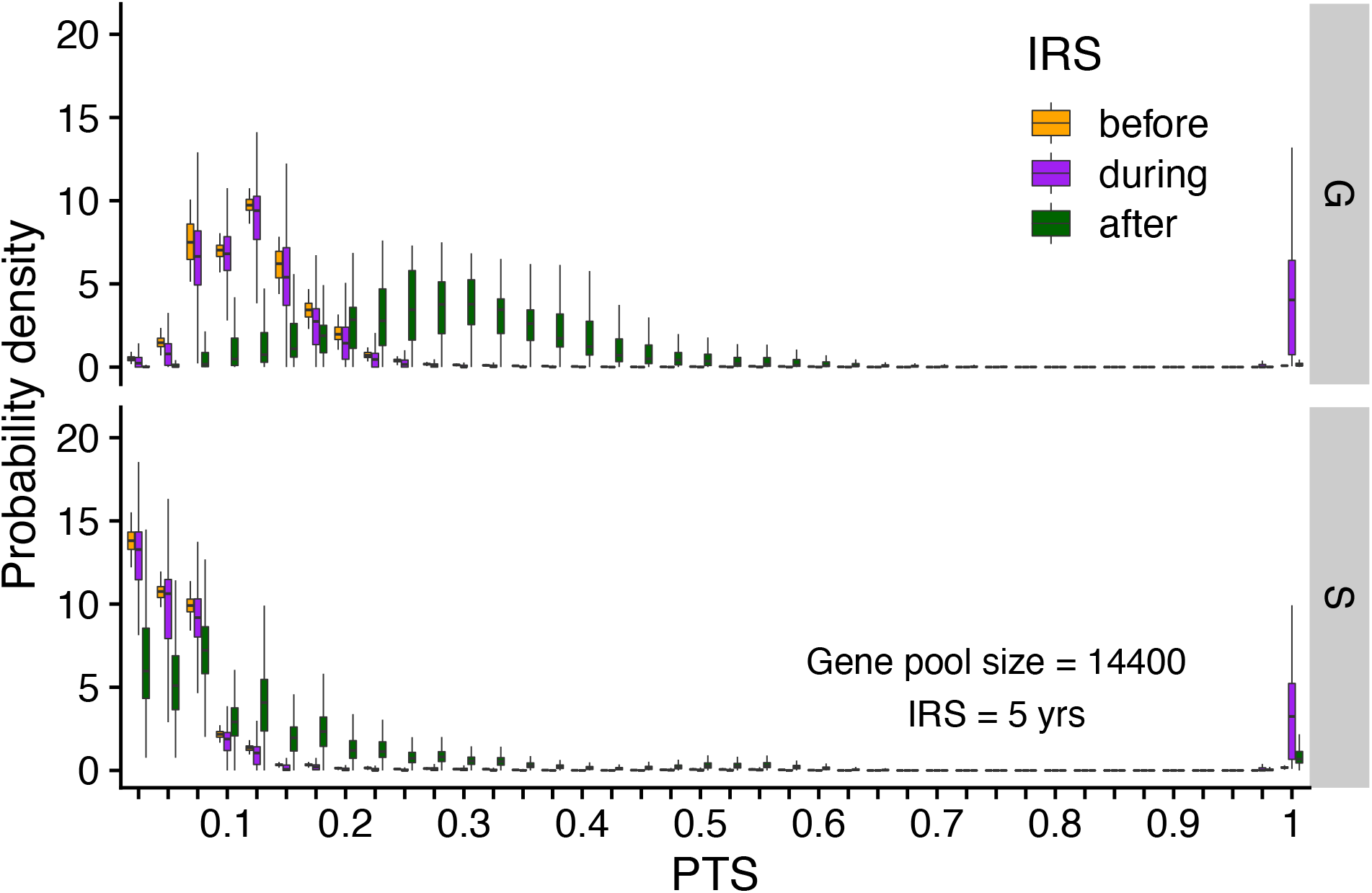
Distribution of pairwise PTS of sampled *var* repertoires before (orange), during (purple) and after (green) interventions between different selection regimes. PTS values are binned into 40 equal sized bins of 0.025. Each box shows the variation of PTS values across different runs. Summarized here are runs from a gene pool size of 14,400, high transmission and a 5-year intervention. The PTS distributions differ between G and S. After intervention, PTS distribution in G drifts away from its initial state whereas in S the distribution tends to go back to its pre-IRS state.

### 3.3 Antigenic diversity recovers independently from prevalence under NFDS

We focus next on the recovery from intervention in terms of both antigenic diversity and prevalence. As expected, both antigenic diversity and prevalence decrease with intervention, and longer IRS leads to a higher reduction in antigenic diversity than 2-year IRS. Nevertheless, diversity recovers more slowly than prevalence (Fig. 5). Despite the slow recovery of antigenic diversity, new genes are strongly preferred under S (partial recovery of Shannon diversity after IRS or even during 10-yr IRS in Fig. 5A), but not under G. All simulations that survive intervention show a comparable initial rebound of prevalence in the months that immediately follow the control period. The rebound for G is to the same prevalence than before intervention as the transmission rate has been restored and the duration of infection only depends on this rate of exposure. In contrast, under S, after an initial overshoot – which is a result of a large human population that is not immune to the parasite due to limited infections during IRS – mean prevalence attains lower values (Fig. S5, Fig. 5B), because the decreased *var* gene diversity induces a shorter duration of infection. The rebound of diversity under S is therefore a result of immune selection and not a larger parasite population size post-IRS. The long-term recovery of prevalence post-intervention can only be achieved slowly given that diversity itself rebuilds slowly.

**Figure 5:**
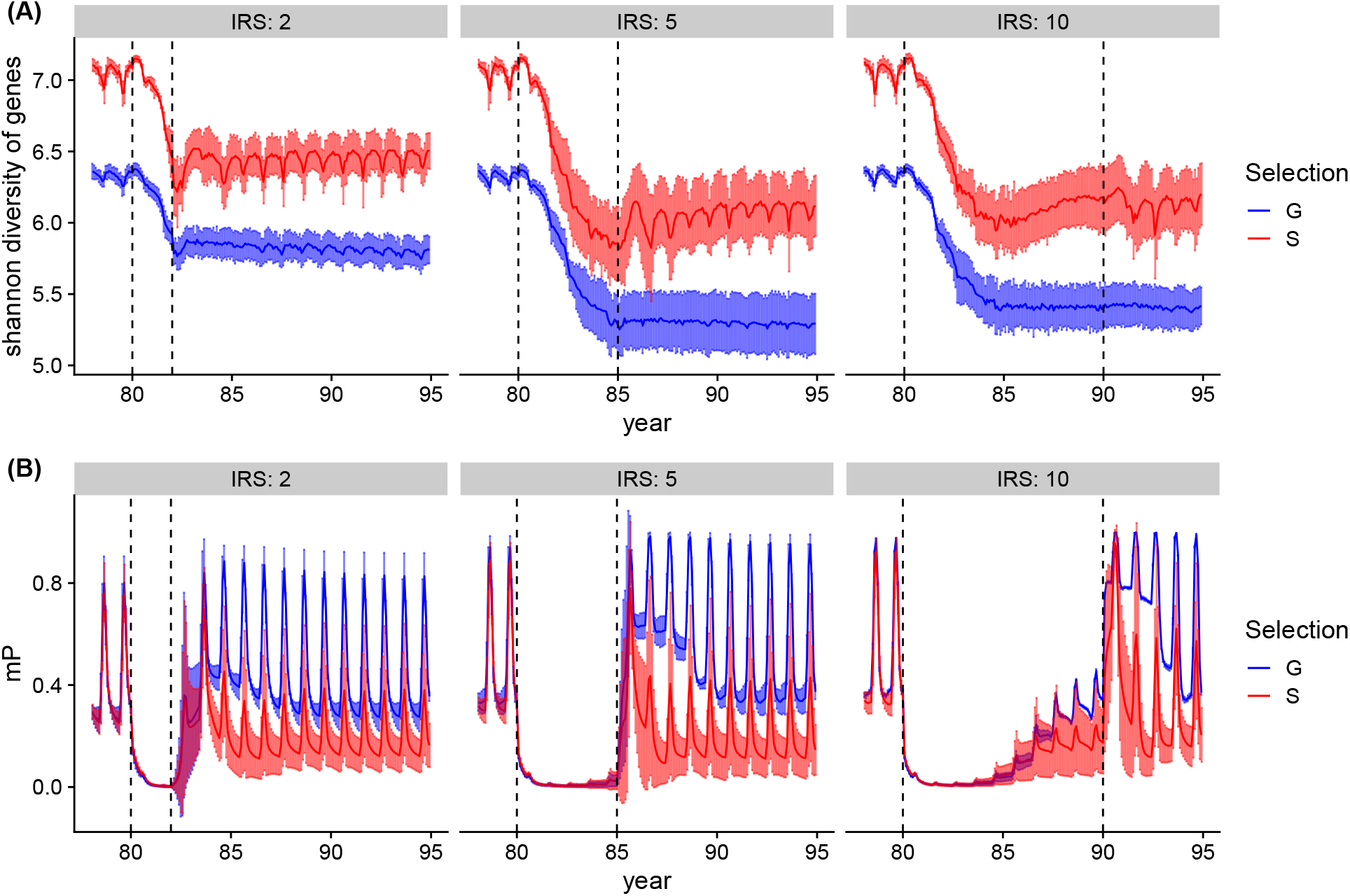
Dynamics of genetic diversity and prevalence during and after IRS. (**A**) Mean Shannon diversity (solid line) and variation (error bars) of *var* gene types is tracked over time. Shaded area shows the extent of the variation across runs. (**B**) Mean changes of prevalence (solid line) and variation (error bars) over time. After IRS, while prevalence under generalized immunity returns back to pre-IRS stage, it gravitates to a lower level under specific immunity. Parameters: gene pool size equals 14,400 and transmission rate is high.

### 3.4 The structure of genetic diversity is restored after intervention under NFDS

Until now we have mainly addressed the connection between antigenic diversity and perturbation in relation to similarity (PTS) as one feature of the structure of diversity. We consider next the structure of similarity between *var* repertoires more broadly by applying an analysis of network features (sensu He *et al*., 2018). In these networks, the nodes are repertoires and the directional links quantify the degree of overlap in *var* gene identity between them. The degree of overlap between two repertoires is a measure of the strength of competition between them for human hosts (i.e., intensity of competition positively correlates with similarity). Therefore, we only retain the links above a given cutoff because it is for these most similar repertoires that we expect the selection against recombinants to be the strongest, and the evidence of frequency-dependent selection to be most apparent (see He *et al*., 2018). We know from previous work that we can use ensembles of network features to differentiate between populations of parasites assembled under S and G. Here, we compare the overall similarity structure between these regimes before, during and after intervention, to ask whether and how quickly the effect of NFDS is restored after intervention, as we expect it to be relaxed under the decreased transmission of intervention.

The similarity structure for an ensemble of simulations can be visualized in the 2-D space defined by the two major axes of variation of PCA (Methods). Figures 6 and S6 show that the networks assembled under S and G do indeed differ and occupy different regions in that space before intervention. Their structure changes and becomes similar for the intervention period as selection/competition is relaxed. Nevertheless, under S the structure is quickly restored after intervention, whereas under G this is not the case, with networks drifting further away from each other and from their initial structure. The network features contributing the most to the classification of the similarity structure are shown in Fig S7. These include in particular the 3-node motifs and reciprocity that characterize local structures and divergence between communities within networks that characterize global structures. The properties distinguish the more tree-like local structure of neutrality from that of limiting similarity and more symmetric or balanced diversity of selection (He *et al*., 2018). The action of frequency-dependent selection on similarity structure is what eventually maintains and restores the distributions of overlap described earlier (Fig 4), which explains the longer duration of infection and therefore the higher probability of persistence during intervention.

**Figure 6:**
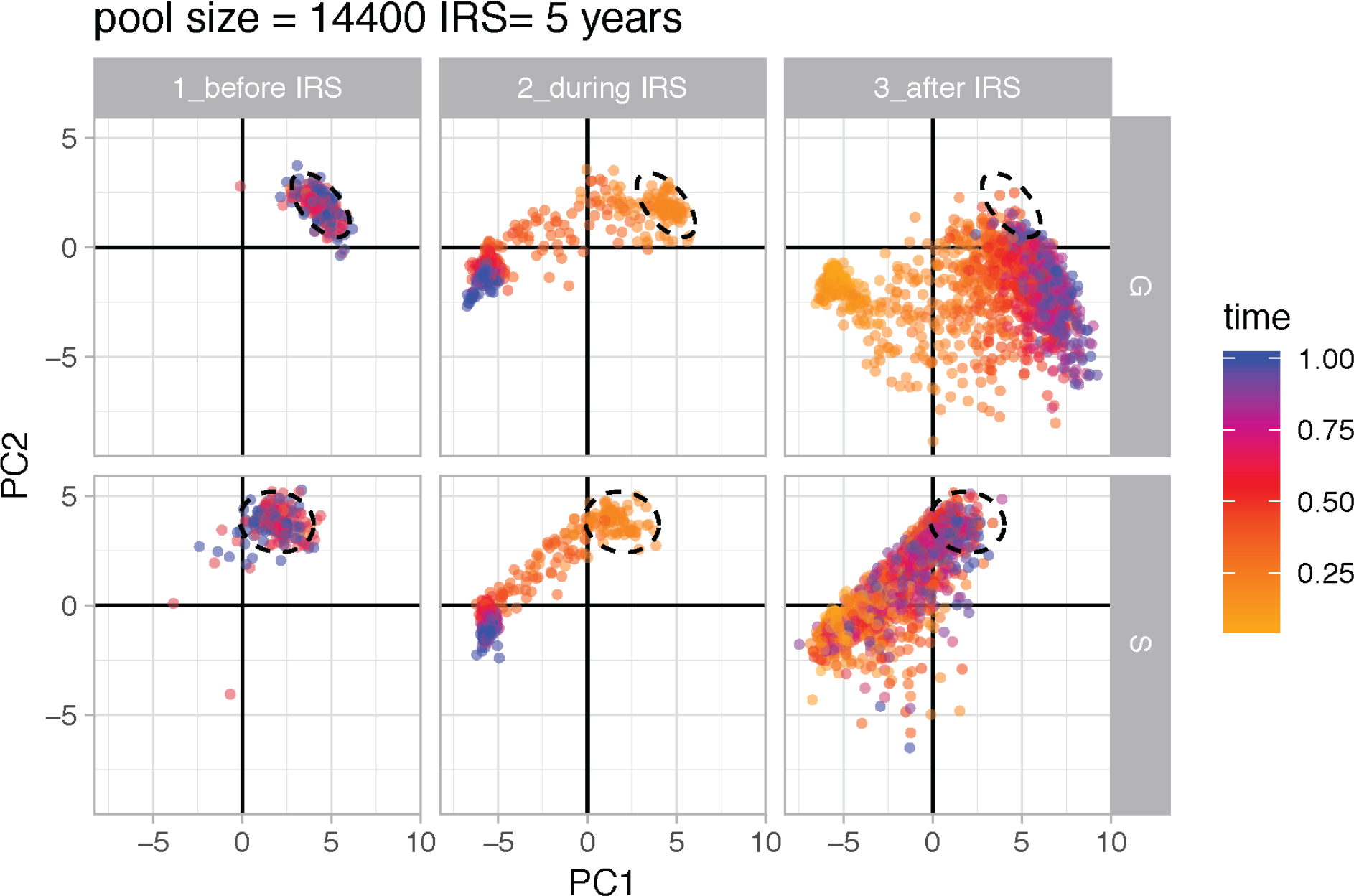
Principal component analysis of 37 network properties of *var* repertoire similarity networks. PCA over time shows how the diversity structures change on the 2D space of the PCs between specific immunity and generalized immunity under high transmission. Colors represent the corresponding time of the network relative to the beginning of the time period. Ellipses show the location of network properties of pre-IRS layers. Parameters: gene pool size equals 14,400 and IRS lasts for 5 years.

## 4 Discussion

Understanding the relationship between structure and resilience in diverse communities has been a long-lasting and on-going effort. This relationship is likely to depend on the processes that generate diversity and assemble its structure. High transmission endemic malaria constitutes a relevant host-pathogen system to investigate this relationship in the context of negative frequency-dependent interactions that stabilize coexistence and structure diversity via patterns of limiting similarity. In the malaria system, limiting similarity emerges from the dominant force behind large antigenic diversity – namely negative frequency-dependent selection from competition of parasites for hosts mediated by adaptive specific immunity. We specifically asked about the connection between the population structure of pathogen genetic diversity and resilience to press perturbations that reduce the abundance of the parasite. We have shown that repertoire populations assembled under selection exerted by host immunity are more resilient to intervention than those assembled under a neutral model with a generalized form of immunity. We have linked this resilience to the larger fraction of highly dissimilar repertoires that provides the parasite with a way to infect non-immune hosts. Interestingly, the network analyses of parasite similarity reveal that the effect of NFDS is quickly restored after the intervention is lifted, which indicates that the process acting to maintain patterns of limiting similarity acts strongly on the parasite population. This quick rebound provides a different angle on the stability of the system. Finally, our results provide an example of how individual and specific interactions determine not only structure, but also stability.

Our theoretical results suggest that competition for hosts can hamper malaria interventions, by operating to maintain the most dissimilar repertoires in tandem with the enormous local diversity (Ruybal-Pesántez *et al*., 2017; Chen *et al*., 2011). This is in line with the challenge of malaria elimination that characterizes high transmission regions with *var* gene diversity level comparable to those considered here, even after intense eradication campaigns. For example, Coleman *et al*. (2017) showed that after lifting a 7-year long IRS effort, transmission intensity (i.e., entomological inoculation rate, EIR) increased from 30 to 90 infectious bites per person per month in only two years, despite a consistent reduction in both EIR and sporozoite rates compared to a nearby control site during the intervention. Thus, transmission persists and prevalence rebounds.

Our model considers a local population embedded within a regional pool that provides a source of genetic variation. This approach is a first step towards developing a more comprehensive theory because in nature malaria is transmitted within and between local human populations, effectively creating a metapopulation of parasites. Therefore, processes that operate on metapopulations such as dispersal and source-sink dynamics, may influence both the assembly of parasite populations and their stability in addition to local selection. For our purpose, this metapopulation context is particularly relevant in creating a vast pool of genetic variation as documented for endemic regions over larger spatial scales (Day *et al*., 2017; Tonkin-Hill, 2020) than those of local transmission, and longer temporal scales than those of the intervention we implement here. Addressing metapopulation dynamics explicitly for this highly diverse system would be however computationally extensive. Our approach relies on the initial compromise of a global pool typical of many assembly models in ecology. We conjecture that our results may apply to other ecological systems where frequency-dependent interactions and resulting selection play an important role in the coexistence of a large number of species. Such interactions arise under stabilizing competition (sensu Chesson, 2000) which generates niche differences, and also under Janzen-Connell mechanisms of negative density-dependent mortality of offspring in the rainforest from interactions with natural enemies and mutualists (Janzen, 1970; LaManna *et al*., 2017; Mangan *et al*., 2010; Schroeder *et al*., 2020). Coexistence under NFDS should apply in particular to systems for which the pool of trait diversity is large, as we have previously argued (Pascual, 2020). Traits of interest associated with negative frequency-dependent mortality in the rainforest concern for example phytochemistry (Forrister *et al*., 2019b). The resulting community structure should in turn increase stability, by reducing overlap between species in trait space, allowing for an overall better ability to cope with perturbations. There is also increasing empirical evidence in other pathogen systems for the importance of frequency-dependent interactions in structuring diversity and determining responses to interventions such as vaccination (Browne & Karubian, 2016; Azarian *et al*., 2019; McNally *et al*., 2019). Our hypothesis remains to be addressed for species communities including through the development of theory. Here, agent-based models and network analyses of the resulting diversity structure should prove useful. Theory can guide network analyses of complex data sets to address the role of non-neutral processes and associated patterns of similarity based on relevant traits.

## 5 Author Contributions

SP, QH and MP conceived the project idea. SP and QH developed simulations and performed computations including network analyses. QH, SP and MP wrote the manuscript. KET and KPD contributed to the model formulation, especially the implementation of the intervention scheme and the migration from a regional pool. All authors discussed the results and contributed to the final manuscript.

## 8 Data Availability Statement

The agent-based stochastic simulator of malaria dynamics is available at the GitHub repository: https://github.com/pascualgroup/varmodel2.

## 9 Supplementary Material

**Table S1:** Epidemiological and genetic parameters used in stochastic simulations.

| Symbol | Type | Description | Values | unit |
| --- | --- | --- | --- | --- |
| $G_p$ | number | initial gene pool size | [1200, 1800, 2400, 3600, 4800, 7200, 9600, 14400, and 19200] | |
| $l$ | number | epitopes | $G_p/10$ | per locus |
| $d$ | time | length of gene expression | 6 | days |
| $\beta$ | rate | contact rate | low = 44<br>medium = 100<br>high = 221 | bites per person per year |
| $\delta$ | rate | immune memory loss rate | 0.001 | per allele per day |
| $m$ | proportion | migration rate | 0.26 | percentage of local transmission |

**Figure S1:**
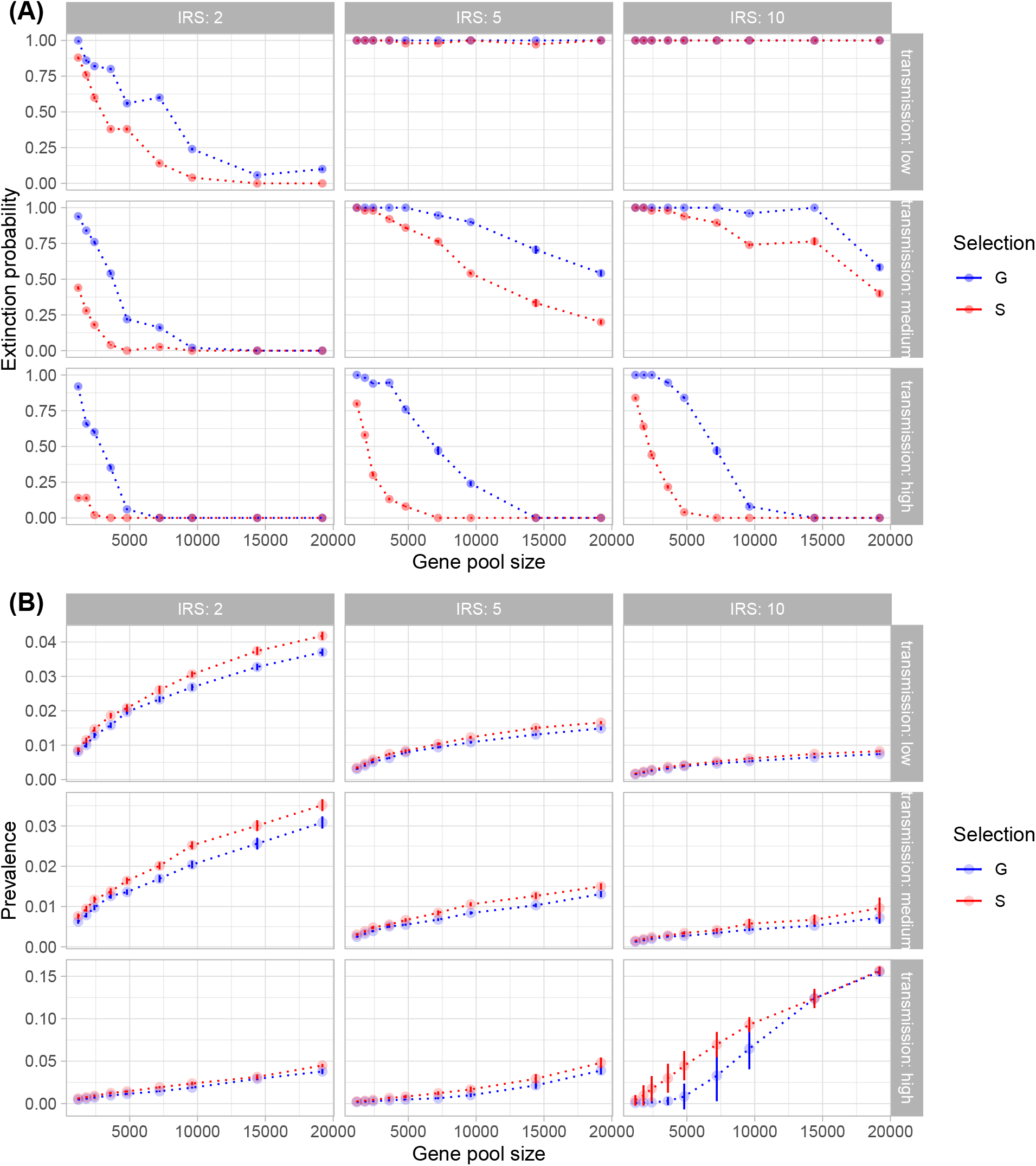
Extinction probability (A) and mean prevalence during IRS (B) as a function of initial gene pool size given under transmission intensities (rows) and lengths of IRS (columns). In (A), each point is the proportion of simulation runs for the given parameter combination (out of 50 runs) that crashed before the IRS was lifted. In (B), each point shows the mean value of 50 replications for the given parameter combination, with bars representing the standard deviation. Blue color depicts simulations under generalized immunity (G, the neutral model), while red color corresponds to specific immunity (S, therefore NFDS).

**Figure S2:**
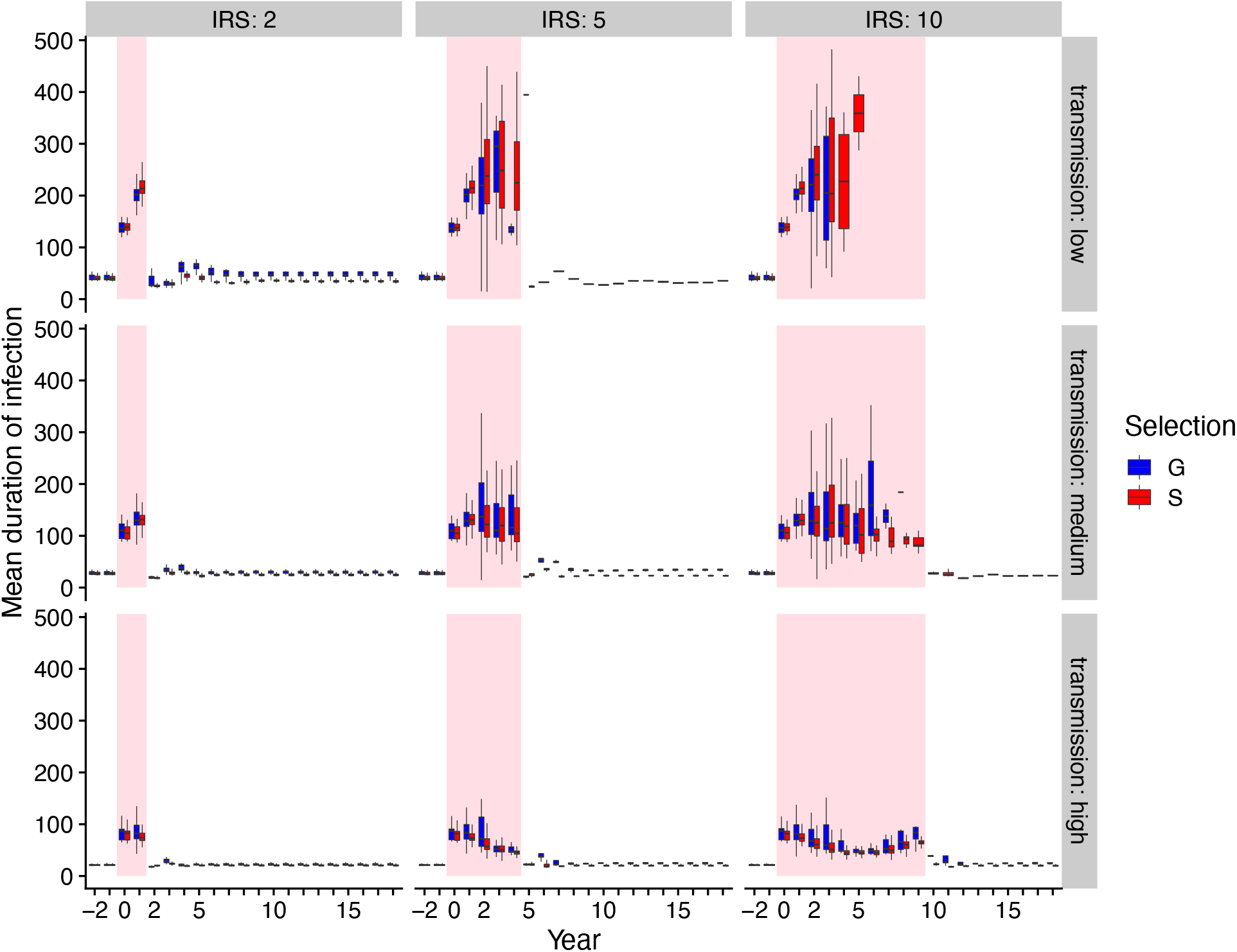
Comparisons of the mean duration of infection over time between S and G given different transmission intensities. Box-plots summarize the distribution of the mean duration of infection across different gene pool sizes and among replication runs. Highlighted areas indicate the IRS period.

**Figure S3:**
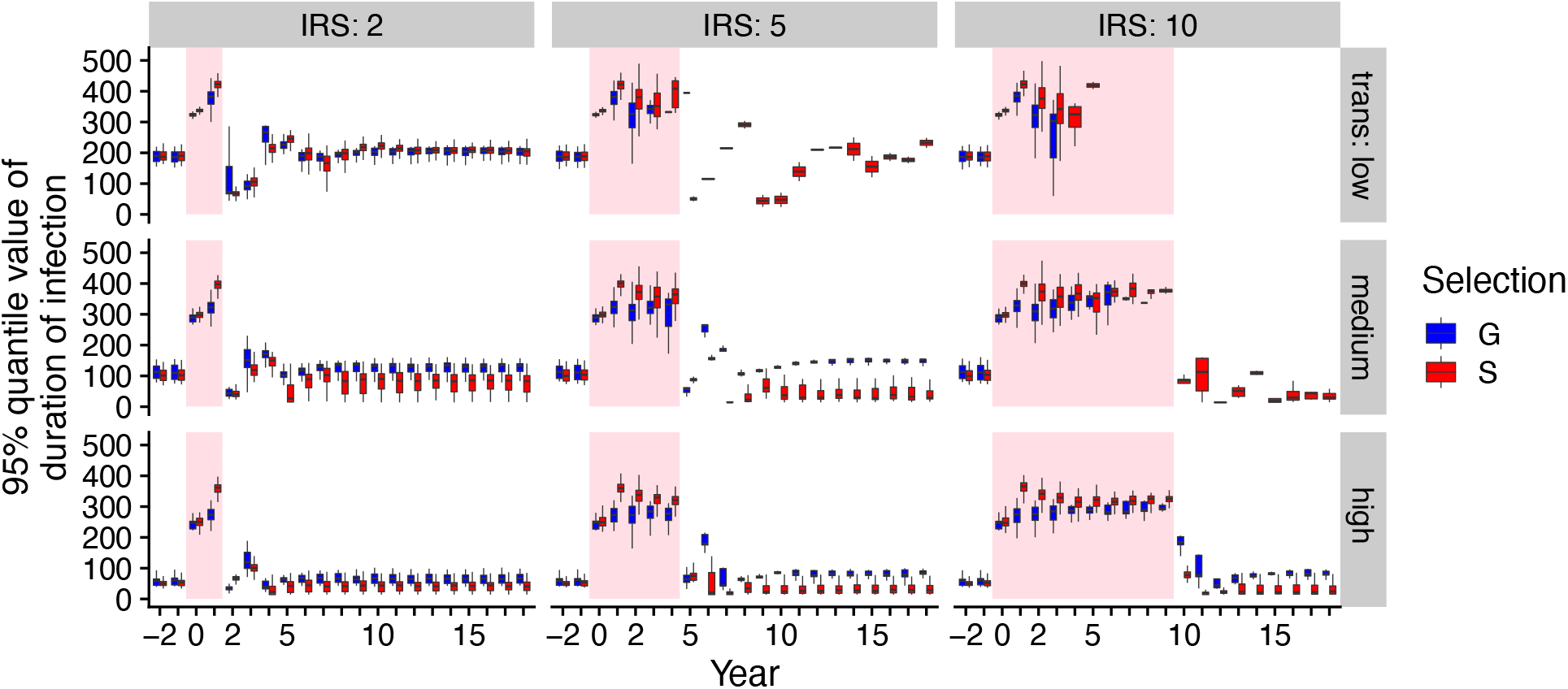
Comparisons of the 95th percentile of duration of infection over time between S and G given different transmission intensities. Boxplot summarizes the distribution of duration of infection in its 95th percentile across different gene pool sizes and among replication runs. Boxes with black frames indicate IRS period.

**Figure S4:**
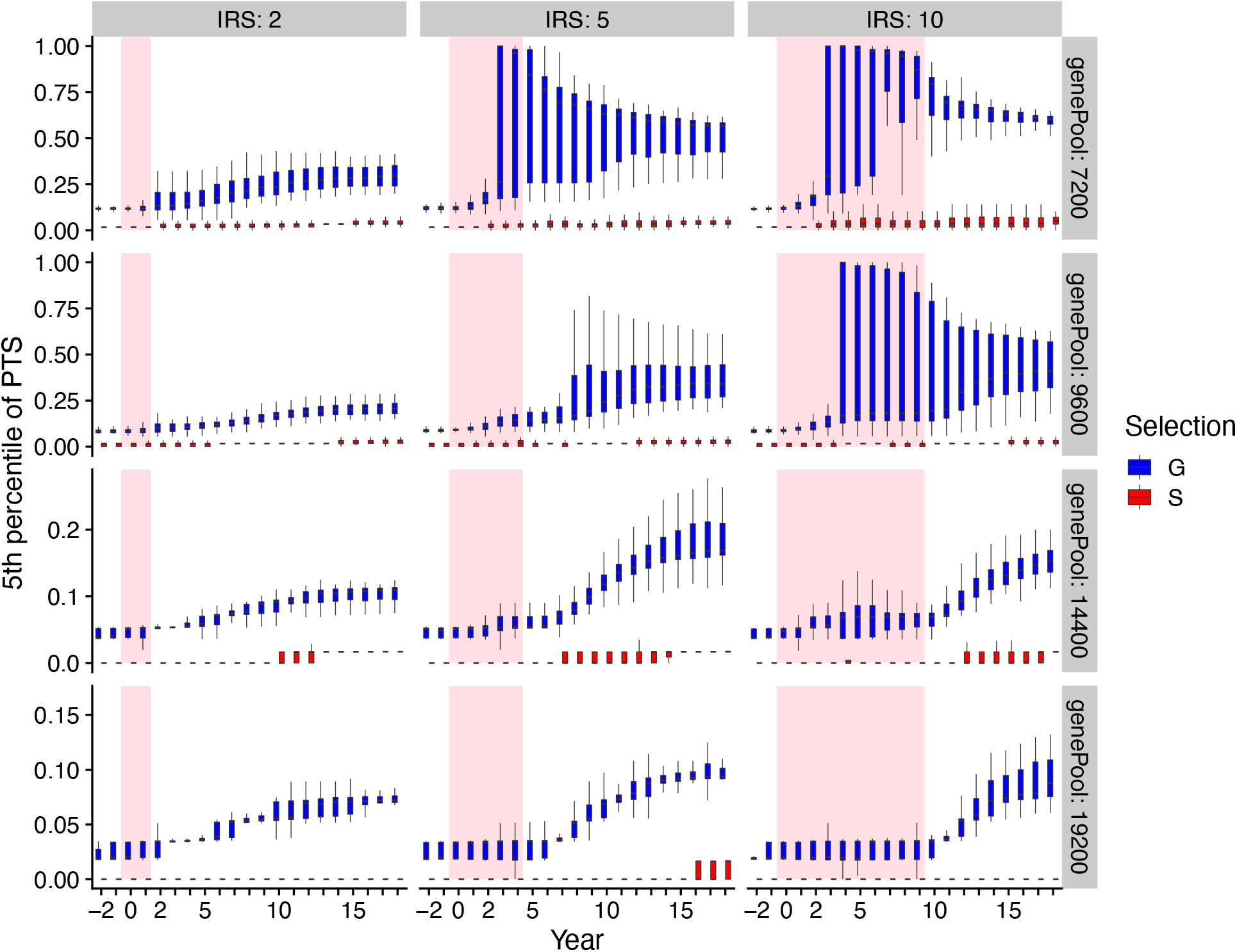
Comparisons of the 5th percentile of PTS over time between between S and G. Box-plots summarize the distribution of PTS in its 5th percentile across different gene pool sizes and among replication runs. Highlighted areas indicate the IRS period.

**Figure S5:**
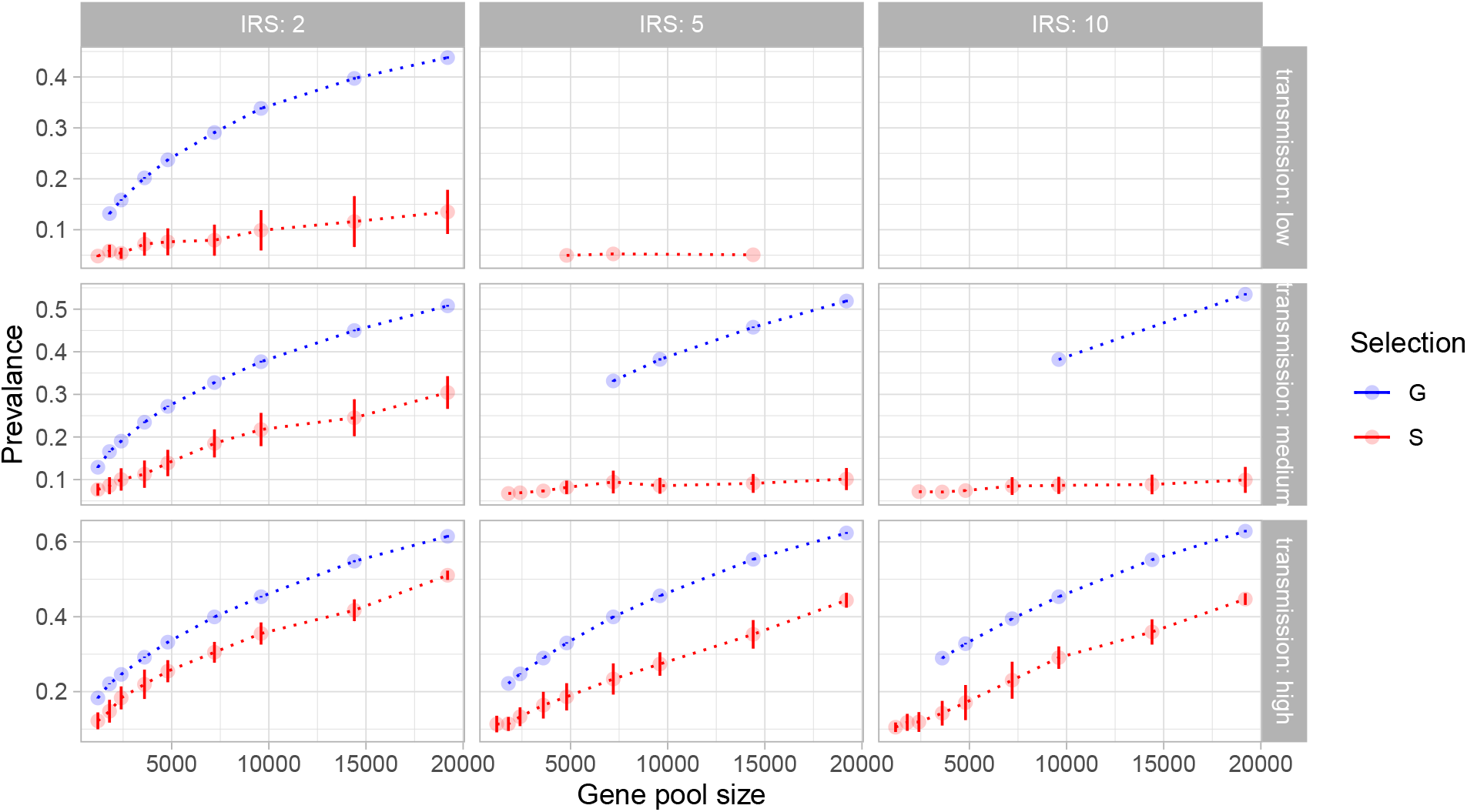
Mean prevalence of surviving runs after IRS as a function of the initial gene pool size given different transmission intensity and length of IRS. Each point shows the mean value (across replications) of the parameter combination with bars representing the standard deviation. Blue color represents simulations under generalized immunity, while red color shows the results from specific immunity.

**Figure S6:**
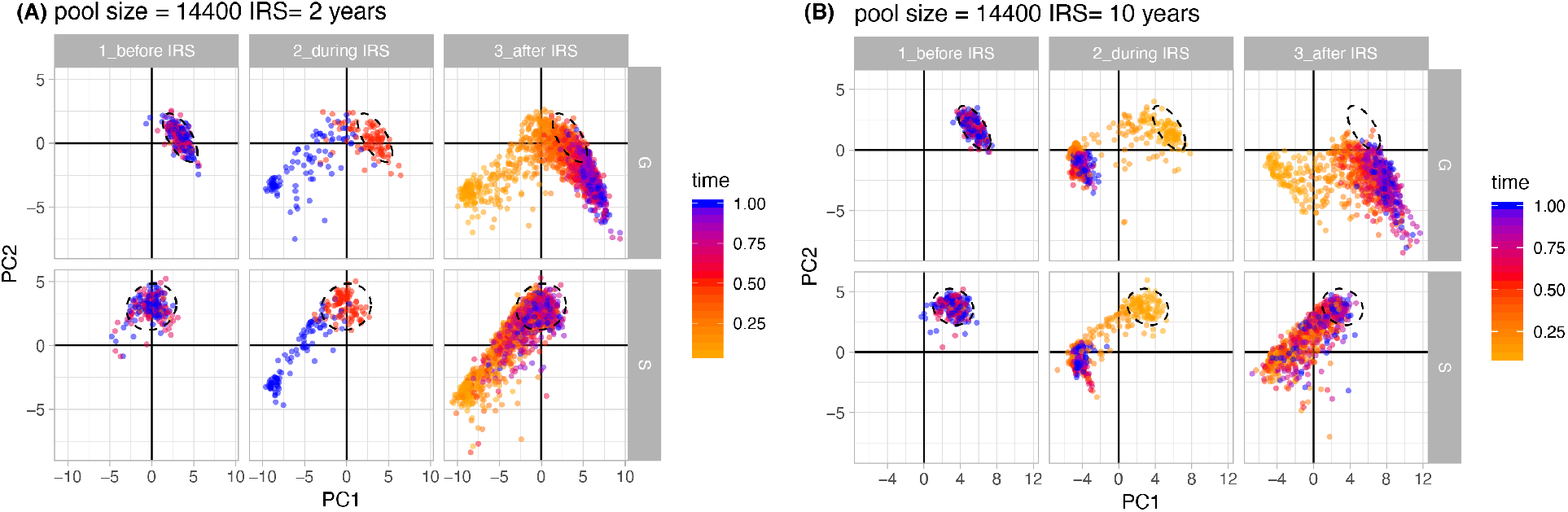
Principal component analysis (PCA) of the dynamics of network properties. Examples of PCA of 37 network properties across time show how the diversity structures change on the 2D space of PCs between specific immunity and generalized immunity under high transmission and high gene pool sizes under 2 and 10 years of interventions. Ellipses show the location of network properties of pre-IRS layers.

**Figure S7:**
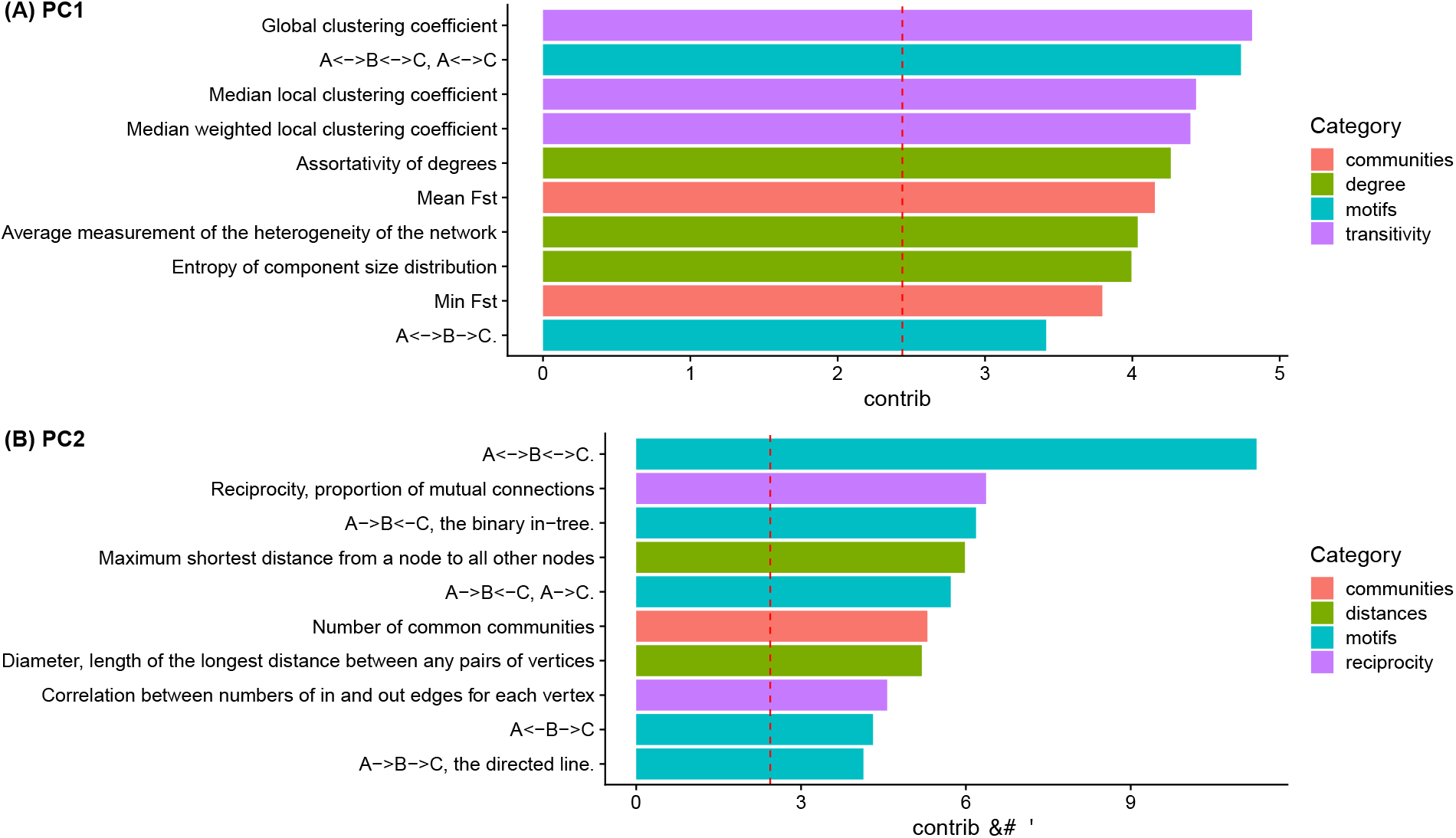
Contributions (in percentage) of network properties to the principal components as shown in Fig. S6. Top 10 properties are shown with a dotted line indicating the expected contribution assuming all variables contribute equally. Color of the bars represents its category of network properties.

